# Reduced long-lasting insecticidal net efficacy and pyrethroid insecticide resistance are associated with over-expression of *CYP6P4, CYP6P3* and *CYP6Z1* in populations of *Anopheles coluzzii* from South-East Côte d’Ivoire

**DOI:** 10.1101/2020.09.24.311639

**Authors:** Anne Meiwald, Emma Clark, Mojca Kristan, Constant Edi, Claire L. Jeffries, Bethanie Pelloquin, Seth R. Irish, Thomas Walker, Louisa A. Messenger

## Abstract

**Background:** Resistance to major public health insecticides in Côte d’Ivoire has intensified and now threatens the long-term effectiveness of malaria vector control interventions.

**Methods:** This study evaluated the bioefficacy of conventional and next-generation long-lasting insecticidal nets (LLINs), determined resistance profiles, and characterized molecular and metabolic mechanisms in wild *Anopheles coluzzii* from South-East Côte d’Ivoire in 2019.

**Results:** Phenotypic resistance was intense: more than 25% of mosquitoes survived exposure to ten times the doses of pyrethroids required to kill susceptible populations. Similarly, 24-hour mortality to deltamethrin-only LLINs was very low and not significantly different to an untreated net. Sub-lethal pyrethroid exposure did not induce significant delayed vector mortality 72 hours later. In contrast, LLINs containing the synergist piperonyl butoxide (PBO), or new insecticides, clothianidin and chlorfenapyr, were highly toxic to *An. coluzzii*. Pyrethroid-susceptible *An. coluzzii* were significantly more likely to be infected with malaria, compared to those that survived insecticidal exposure. Pyrethroid resistance was associated with significant over-expression of *CYP6P4, CPY6Z1* and *CYP6P3*.

**Conclusions:** Study findings raise concerns regarding the operational failure of standard LLINs and support the urgent deployment of vector control interventions incorporating PBO, chlorfenapyr or clothianidin in areas of high resistance intensity in Côte d’Ivoire.

## Introduction

In Côte d’Ivoire, malaria is a serious public health problem with the entire population of ∼26.2 million people is at risk, and disease prevalence reaching as high as 63% in the south-west region [1]. Control of *Anopheles gambiae* s.l., the major malaria vector species group in Côte d’Ivoire, has been through the efforts of the National Malaria Control Programme (NMCP), which has distributed insecticide-treated nets (ITNs) as the primary vector control intervention. Indoor residual spraying (IRS) and larviciding in high transmission areas have been recommended as complementary strategies; implementation of the former has commenced in late 2020 [2]. Estimates of net coverage across the country remain low, with the proportion of households with at least one ITN for every two people rising from 31% in 2012 to 47% in 2016, and ITN use stagnating at 40% of households reporting sleeping under a net the previous night in both survey years [2]. The most recent universal net campaigns in Côte d’Ivoire in 2017–2018 issued conventional, pyrethroid (deltamethrin) long-lasting insecticidal nets (LLINs), aiming to achieve 90% coverage and 80% use [2]. However, country-wide, multi-class insecticide resistance among populations of *An. gambiae* s.l. is a growing cause for concern because of potential operational failure of current vector control strategies, both locally, as well as across the sub-Saharan region [2,3].

Resistance to pyrethroid and carbamate insecticides in *Anopheles* mosquitoes was first reported from the central region of Côte d’Ivoire in the early 1990s [4-7]. Subsequently, local resistance to the major insecticide classes recommended by the World Health Organization (WHO) for adult mosquito control – pyrethroids, carbamates, organophosphates, and organochlorines – evolved rapidly [8–10] and has been increasing in intensity, driven largely by selective pressures imposed by contemporaneous scale-up of public health vector control interventions (including those targeting malaria, trypanosomiasis and onchocerciasis vectors) and use of agricultural pesticides [7, 11–14]. This escalation in resistance has now begun to compromise the insecticidal efficacy and community-wide impact of conventional, pyrethroid LLINs in Côte d’Ivoire [14,15], although some levels of personal protection may still remain [15–17].

Amongst vector populations across Côte d’Ivoire, the L1014F *kdr* mutation is pervasive and has been implicated in some longitudinal trends in decreasing DDT and pyrethroid susceptibility [7, 11]; L1014S *kdr* and N1575Y resistance mutations have also been detected but at much lower frequencies [18]. Extreme carbamate (bendiocarb) resistance and pyrethroid cross-resistance in local *An. gambiae* s.s. populations have been shown to be mediated by over-expression of *CYP6P3* and *CYP6M2* and duplication of the G119S *Ace-1* mutation [19].

To support and safeguard future malaria control efforts in Côte d’Ivoire, this study evaluated the efficacy of conventional and next-generation LLINs for prospective distribution; determined current insecticide resistance profiles of *An. gambiae* s.l. (principally *An. coluzzii*); and characterized underlying molecular and metabolic resistance mechanisms.

## Methods

### Study area and mosquito collections

The study protocol was approved by the Comité National d’Ethique des Sciences de la Vie et de la Santé (#069-19/MSHP/CNESVS-kp) and the London School of Hygiene and Tropical Medicine (#16782 and #16899). Study activities were conducted in the village of Aboudé, rural Agboville, Agnéby-Tiassa region, south-east Côte d’Ivoire (5^°^55’N and 4^°^13’W), selected due to its high mosquito densities and malaria prevalence (26% in children <5 years old, in recent estimates [1]). Adult mosquitoes were collected nightly between 5^th^ July and 26^th^ July 2019, using human landing catches (HLCs), inside and outside households from 18:00 to 06:00hr. Unfed mosquitoes, morphologically identified as *An. gambiae* s.l. [20], were tested in bioassays that same day, following a brief recovery period; blood-fed mosquitoes were first held for 2–3 days to allow for blood-meal digestion.

### WHO cone bioassay testing

Two types of LLIN were evaluated in this study. PermaNet^**®**^ 2.0 is a conventional LLIN treated with deltamethrin only (1.4g/kg±25%) and PermaNet^**®**^ 3.0 is a PBO synergist LLIN, consisting of a roof containing PBO (25g/kg) and deltamethrin (4g/kg±25%) and side panels containing deltamethrin only (2.8g/kg±25%). WHO cone bioassays were used to test the susceptibility of *An. gambiae* s.l. exposed to unwashed PermaNet^**®**^ 2.0, PermaNet^**®**^ 3.0 roof panels and PermaNet^**®**^ 3.0 side panels [21]. To control for potential variation in insecticide/synergist content, each of five LLINs per type was cut into 19 pieces, measuring 30 × 30cm, with each piece tested a maximum of three times.

### Resistance intensity and synergist bioassay testing

Centers for Disease Control and Prevention (CDC) resistance intensity bioassays were performed for six public health insecticides (pyrethroids: alpha-cypermethrin, deltamethrin and permethrin; carbamate: bendiocarb; neonicotinoid: clothianidin; and pyrrole; chlorfenapyr) [22,23]. The diagnostic doses of all insecticides were evaluated (including clothianidin: 90µg/bottle [23] and chlorfenapyr: 100µg/bottle) and 2, 5 and 10 times the diagnostic dose of pyrethroid insecticides were also used. Per test, knock-down was recorded at 15-minute intervals for 30 minutes (pyrethroids and bendiocarb) or 60 minutes (clothianidin and chlorfenapyr) of insecticide exposure. PBO pre-exposures were performed using WHO tube assays [24], prior to CDC bottle bioassay testing.

WHO cone and CDC resistance intensity bioassay data were interpreted according to the WHO criteria [21,22]. Mosquitoes which died following exposure to a LLIN or 1X insecticide dose were stored at −20°C in RNAlater^®^ (Thermo Fisher Scientific, UK) and were considered ‘susceptible’ for genotypic analysis. Surviving mosquitoes were held and scored for mortality after 24, 48 and 72 hours to observe delayed mortality. Kaplan-Meier curves were used to visualize survival data, and Cox regression was used to compare post-exposure survival. Immediate mortality following LLIN (60 minutes and 24 hours) or insecticidal exposure (30 or 60 minutes, depending on insecticide) were excluded. Surviving mosquitoes at 72 hours were stored at −20°C in RNAlater^®^ and were considered ‘resistant’ for genotypic analysis.

### Mosquito processing, identification of *Anopheles gambiae* s.l. species complex members and *Plasmodium falciparum* detection

A sub-sample of field-caught mosquitoes that were tested in bioassays were selected for molecular analysis (n=912). Approximately equal numbers of specimens were chosen to represent phenotypically ‘susceptible’ or ‘resistant’ mosquitoes for each LLIN type or insecticide dose, and selected across different replicates/testing days to capture as much population-level variation as possible. RNA was extracted from individual whole-body mosquitoes according to standard protocols [23]. Field *An. gambiae* s.l. were identified to species-level by amplification of the SINE200 insertion that differentiates *An. coluzzii* and *An. gambiae* s.s. [25] and were screened for the presence of *Plasmodium falciparum* [26].

### Characterization of insecticide resistance mechanisms: target site mutations

The same cohort of field mosquitoes (n=912) were tested for the presence of the L1014F *kdr* [27] and N1575Y mutations [28]. A sub-sample of mosquitoes (n=49) which were exposed to bendiocarb, clothianidin or chlorfenapyr were tested for the presence of the G119S *Ace-1* mutation [29]. Pearson’s Chi-squared tests and Fisher’s exact tests (when sample sizes were small) were used to investigate the statistical association between resistance status, allele frequencies and deviations from Hardy-Weinberg equilibrium.

### Characterization of insecticide resistance mechanisms: metabolic gene expression

Relative expression of five metabolic genes (*CYP6P3, CYP6P4, CYP6Z1 CYP6P1* and *GSTE2*) was measured in all field collected mosquitoes (n=912), using multiplex quantitative real-time PCR (qRT-PCR) assays, relative to the housekeeping gene ribosomal protein S7 (*RPS7*) [30]. In addition, gene expression levels were measured in susceptible *An. coluzzii* N’gousso colony mosquitoes (n=48). All samples were run in technical triplicate. Relative expression level and Fold Change (FC) of each target gene from resistant and susceptible field samples, relative to the susceptible laboratory strain, were calculated using the 2^-ΔΔCT^ method incorporating PCR efficiency, normalised relative to the endogenous control gene (*RPS7*).

## Results

### Mosquito collections and species identification

A total of 4,609 female *An. gambiae* s.l. mosquitoes were collected in Agboville, Côte d’Ivoire. Of those, 912, which were previously tested in either LLIN bioefficacy assays (n=384) or resistance intensity bioassays (n=528), were selected for molecular species identification, with 805 (88.3%) determined to be *An. coluzzii*, 75 (8.2%) *An. gambiae* s.s. and 22 (2.4%) *An. gambiae*-*An. coluzzii* hybrids; 10 individuals did not amplify.

### Long-lasting insecticidal net efficacy

A total of 2,666 field-caught *An. gambiae* s.l. were used to assess the bioefficacy of conventional pyrethroid-treated LLINs (PermaNet^**®**^ 2.0 and PermaNet^**®**^ 3.0 side panels) and next-generation synergist LLINs (PermaNet^**®**^ 3.0 roof panels), compared to an untreated control (Figure 1).

**Figure 1.**
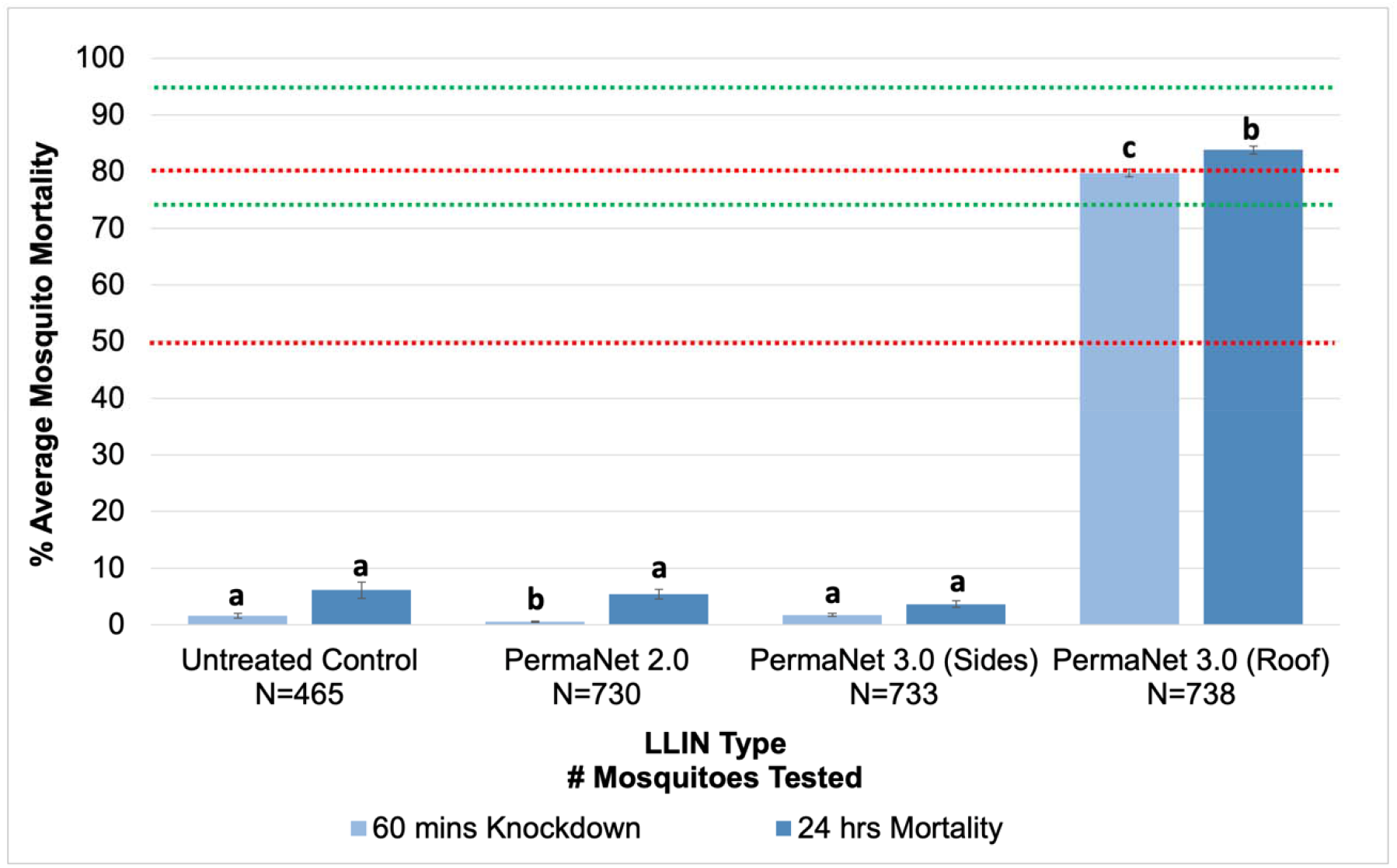
Bioefficacy of different unwashed LLINs against field-caught *An. gambiae* s.l. Mean knock-down and mortality rates with 95% confidence intervals (CI) at 60 minutes and 24 hours, respectively, after 3 minutes exposure to PermaNet**®** 2.0 (deltamethrin only), side panels of PermaNet**®** 3.0 (deltamethrin only), roof panels of PermaNet**®** 3.0 (PBO + deltamethrin) and an untreated control net. Knock-down or mortality in the same time period for each treatment sharing a letter do not differ significantly (*p*>0.05). Green lines at ≥75% knock-down = minimal effectiveness at 60 minutes and at ≥ 95% knock-down = optimal effectiveness at 60 minutes. Red lines at ≥50% mortality = minimal LLIN effectiveness at 24 hours and ≥80% mortality = optimal LLIN effectiveness at 24 hours, as defined by the WHO [21].

Overall, levels of *An. gambiae* s.l. knock-down and mortality to deltamethrin LLINs, were very low and largely equivalent to the untreated control net (Figure 1). At 60 minutes, average mosquito knock-down to the untreated control, PermaNet^**®**^ 2.0 and PermaNet^**®**^ 3.0 side panels was 1.56% (95% CI: 1.13-1.99%), 0.54% (95% CI: 0.42-0.65%) and 1.75% (95% CI: 1.49-2.0%), respectively. By contrast, average mosquito knock-down for PBO-containing PermaNet^**®**^ 3.0 roof panels was significantly higher (79.8%, 95% CI: 79.07-80.48%; χ^2^ =705.51, 968.65 and 937.33; *p*<0.001, *versus* untreated control, PermaNet^**®**^ 2.0 and PermaNet^**®**^ 3.0 side panels, respectively) (Figure 1).

At 24 hours, mortality to the untreated control, PermaNet^**®**^ 2.0 and PermaNet^**®**^ 3.0 side panels remained low (6.11%, 95% CI: 4.71-7.51%; 5.44%, 95% CI: 4.58-6.29% and 3.66%, 95% CI: 3.12-4.19%, respectively), while mortality to PermaNet^**®**^ 3.0 roof panels increased only marginally but still remained significantly higher (83.81%, 95% CI: 83.15-84.47%; χ^2^ =727.96, 914.61 and 963.09; *p*<0.001 for all, *versus* untreated control, PermaNet^**®**^ 2.0 and PermaNet^**®**^ 3.0 side panels, respectively) (Figure 1). PermaNet^**®**^ 3.0 roof panels reached minimal effectiveness (knock-down ≥75%) 60 minutes after exposure and optimal effectiveness (mortality ≥80%) at 24 hours. Neither of the deltamethrin-only LLINs reached either effectiveness threshold at any time point.

### Insecticide resistance intensity

One thousand, nine hundred and forty-three field-caught *An. gambiae* s.l. were tested in resistance bioassays. Intense pyrethroid resistance was evident with more than 25% of mosquitoes surviving exposure to ten times the dose of insecticide required to kill a susceptible population; at the diagnostic dose, mosquito mortality did not exceed 25% for any pyrethroid tested (Figure 2A). These results are consistent with the high survival rates observed during cone bioassays using conventional LLINs (Figure 1). In general, levels of resistance to alpha-cypermethrin, deltamethrin and permethrin were not significantly different at each insecticide concentration tested (Figure 2A).

**Figure 2.**
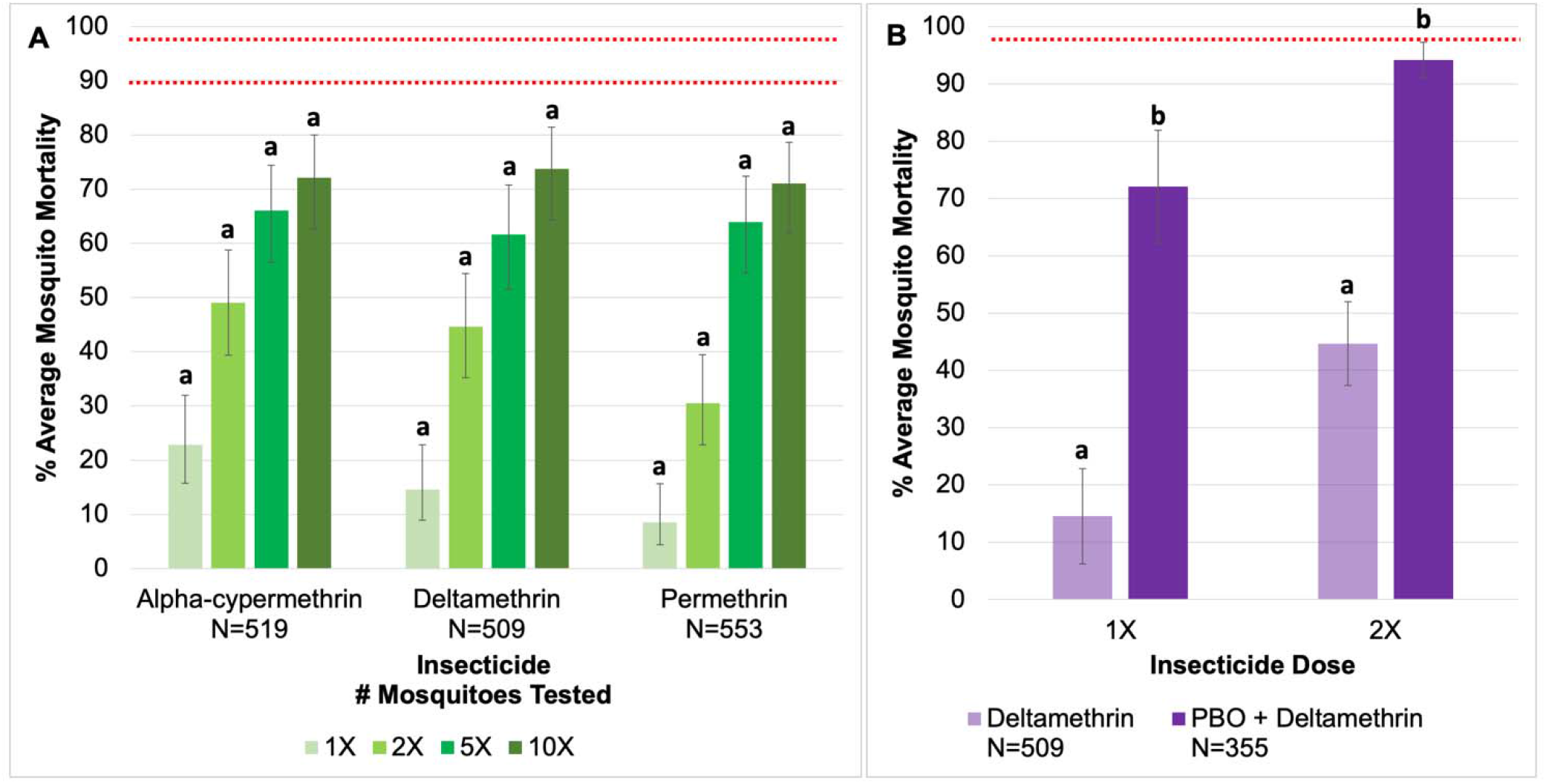
**A:** Resistance intensity of field-caught *An. gambiae* s.l. after exposure to one, two, five and ten times the diagnostic dose of pyrethroid insecticides. Mean knock-down/acute toxicity after 30 minutes exposure with 95% confidence intervals (CI). Knock-down/mortality at the same dose per insecticide sharing a letter do not differ significantly (*p*>0.05). Values of less than 90% mortality (lower red line) indicate confirmed resistance at the diagnostic dose (1X) and values of less than 98% mortality (upper red line) indicate moderate to high intensity resistance or high intensity resistance at 5X and 10X, respectively, as defined by the WHO [24]. **B:** Restoration of deltamethrin susceptibility of field-caught *An. gambiae* s.l. after pre-exposure to PBO. Mean knock-down/acute toxicity after 30 minutes exposure to one or two times the diagnostic dose of deltamethrin with 95% confidence intervals (CI). Knock-down/mortality between pyrethroid only and synergist + pyrethroid sharing a letter do not differ significantly (*p*>0.05). Red line at 98% mortality indicates metabolic resistance mechanisms partially involved [24].

By comparison, carbamate tolerance was low, with mean knock-down of 94.53% (95% CI: 92.11-96.95%) after 30 minutes exposure to the diagnostic dose of bendiocarb. Similarly, high levels of susceptibility to new insecticides clothianidin and chlorfenapyr were observed, with mean mortality of 94.11% (95% CI: 93.43-94.80%; n=102) and 95.54% (95% CI: 94.71-96.36%; n=112), respectively, 72 hours after exposure to the tentative diagnostic doses.

Pre-exposure to PBO increased average *An. gambiae* s.l. mortality significantly from 14.56% (95% CI: 6.24-22.88%) to 72.73% (95% CI: 64.81-79.43) and from 44.66% (95% CI: 34.86-54.46%) to 94.17% (95% CI: 91.12-97.22) after exposure to one or two times the diagnostic dose of deltamethrin (Figure 2B).

### Mosquito survival following insecticidal exposure

All *An. gambiae* s.l. tested in LLIN bioefficacy or resistance intensity bioassays, were held for 72 hours, to assess any impact of insecticide or net exposure on delayed mortality. For LLIN bioassays, there was little evidence for any reduction in survival during this holding period (Cox regression *P*= 0.149, 0.272 and 0.85 comparing PermaNet^**®**^ 2.0, PermaNet^**®**^ 3.0 side panels and PermaNet^**®**^ 3.0 roof panels *versus* untreated control, respectively) (Table 1 and Figure 3A). Exposure to the diagnostic doses of all insecticides in CDC bottle bioassays did not induce significant delayed mortality over 72 hours (Cox regression *P*>0.05 for all insecticides compared to the control; with the exception of chlorfenapyr, *P*=0.02) (Table 1 and Figure 3B). This phenomenon was also observed at increasing pyrethroid doses (Cox regression *P*>0.05 for alpha-cypermethrin, deltamethrin and permethrin 5X and 10X *versus* either the control or diagnostic dose) (Table 1; Figure 3C and 3D).

**Table 1.** Cox proportional hazard model to describe the impact of LLIN/insecticidal exposure on survival of field-caught *An. gambiae* s.l. 72 hours post exposure.

| Insecticide Exposure | N (N Events) | HRR | 95% CI | P-value |
| --- | --- | --- | --- | --- |
| Untreated Netting |  | Reference | - | - |
| PermaNet® 2.0 (deltamethrin only) | 1135 (1047) | 1.095 | 0.968-1.239 | 0.149 |
| PermaNet® 3.0 side panels (deltamethrin only) | 1157 (1088) | 0.9664 | 0.9092-1.027 | 0.272 |
| PermaNet® 3.0 roof panels (PBO + deltamethrin) | 563 (533) | 1.007 | 0.939-1.079 | 0.85 |
| Acetone Control |  | Reference | - | - |
| Alpha-cypermethrin 1X | 676 (641) | 1.006 | 0.9696-1.043 | 0.767 |
| Deltamethrin 1X | 683 (645) | 0.9942 | 0.9539-1.036 | 0.782 |
| Permethrin 1X | 693 (661) | 1.015 | 0.9698-1.062 | 0.525 |
| Clothianidin 1X | 698 (581) | 1.208 | 0.9227-1.581 | 0.169 |
| Chlorfenapyr 1X | 708 (580) | 1.692 | 1.086-2.637 | **0.02** |
| PBO + Deltamethrin 1X | 630 (577) | 0.9662 | 0.2411-3.873 | 0.961 |
| Alpha-cypermethrin 5X | 633 (601) | 0.9951 | 0.9407-1.053 | 0.863 |
| Deltamethrin 5X | 652 (610) | 0.9942 | 0.9393-1.052 | 0.842 |
| Permethrin 5X | 636 (583) | 0.9931 | 0.8638-1.142 | 0.923 |
| Alpha-cypermethrin 10X | 624 (587) | 0.9951 | 0.917-1.08 | 0.906 |
| Deltamethrin 10X | 623 (588) | 0.9943 | 0.9072-1.09 | 0.902 |
| Permethrin 10X | 656 (603) | 1.026 | 0.9509-1.107 | 0.509 |
| 1X Insecticide Dose |  | Reference | - | - |
| Alpha-cypermethrin 5X | 117 (92) | 1.016 | 0.9069-1.138 | 0.785 |
| Alpha-cypermethrin 10X | 108 (78) | 1.007 | 0.9403-1.078 | 0.845 |
| Deltamethrin 5X | 143 (105) | 1.0 | 0.9035-1.107 | 1.0 |
| Deltamethrin 10X | 114 (83) | 1.0 | 0.9363-1.068 | 1.0 |
| Permethrin 5X | 137 (94) | 1.022 | 0.8528-1.225 | 0.812 |
| Permethrin 10X | 157 (114) | 0.9952 | 0.9491-1.044 | 0.842 |

HRR: hazard rate ratio; ratio between the hazard rate in control/reference group and hazard rate for each treatment group.
Significance level defined as $\alpha = 0.05$ .

**Figure 3.**
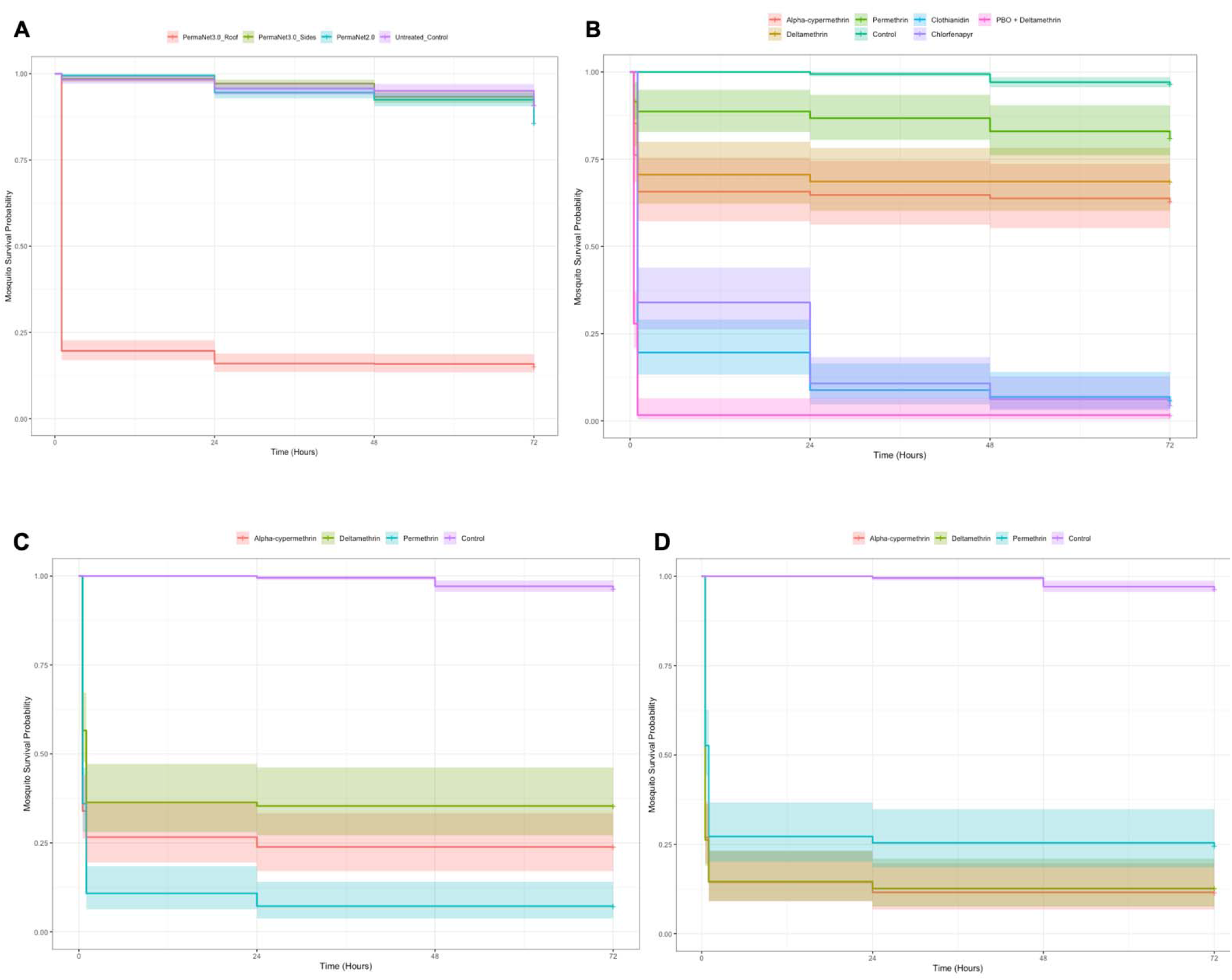
The longevity of field-caught *An. gambiae* s.l. after exposure to LLINs in WHO cone assays (**A**) 1X (**B**), 5X (**C**) and 10X (**D**) times the diagnostic dose of pyrethroid insecticides in CDC resistance intensity assays. Kaplan Meier survival curves indicate the proportion alive each day post-exposure. Immediate mortality following LLIN (60 minutes and 24 hours) or insecticidal exposure (30 or 60 minutes, insecticide depending) were excluded.

### Malaria prevalence

Of the 912 *An. gambiae* s.l. mosquitoes assayed, 31 tested positive for *P. falciparum* (3.4%). For PCR-confirmed *An. coluzzii, P. falciparum* prevalence was 3.50% (28/805); the remaining three infections were in *An. gambiae* s.s. (4%; 3/75). By resistance phenotype, susceptible *An. coluzzii* (i.e. those which died following pyrethroid exposure) were more likely to be infected with malaria, compared to resistant mosquitoes (□^2^ =4.6987; *p*=0.030); infection rates were 5.94% (13/219) and 2.49% (10/401), respectively.

### Target site resistance mutations

L1014F *kdr* screening revealed 92.2% (796/863) of *An. gambiae* s.l. mosquitoes harboured the mutation; 71.5% (617/863) were homozygous, 20.7% (179/863) were heterozygous, 5.1% (44/863) were wild type and 2.6% (23/863) did not amplify. For PCR-confirmed *An. coluzzii*, L1014F *kdr* prevalence was 87.8% (707/805); 66.6% (536/805) were homozygous for the mutation, 21.2% (171/805) were heterozygous, 5.3% (43/805) were wild type and 2.2% (18/805) did not amplify. For *An. coluzzii*, population-level L1014F *kdr* allele frequency was 0.83, with evidence for significant deviations from Hardy-Weinberg equilibrium (□^2^ =29.124; *p*<0.0001). There was no significant association between L1014F *kdr* frequency and ability of mosquitoes to survive pyrethroid exposure, in either LLIN or resistance bioassays (□^2^ =2.0001; *p*=0.157 and □^2^ =3.7577; *p*=0.0.53, respectively). Similarly, there was no significant association between L1014F *kdr* and ability of mosquitoes to survive PBO pre-exposure and pyrethroid treatment, in either LLIN or resistance bioassays (□^2^ =0.0086; *p*=0.926, Fisher’s exact=0.429, respectively). For PCR-confirmed *An. gambiae* s.s., L1014F *kdr* prevalence was 95.3% (61/64); 89.1% (57/64) were homozygous for the mutation, 6.3% (4/64) were heterozygous, none were wild type and 4.7% (3/64) did not amplify. For *An. gambiae* s.s., population-level L1014F *kdr* allele frequency was 0.97, with no significant deviations from Hardy-Weinberg equilibrium (□^2^ =0.070; *p*=0.791).

N1575Y screening revealed 2.3% (21/912) of *An. gambiae* s.l. mosquitoes harboured the mutation; all of these were heterozygotes. N1575Y prevalence was 1.1% (9/805) and 16% (12/75) for PCR-confirmed *An. coluzzii* and *An. gambiae* s.s., respectively; 0.99% (9/912) did not amplify. There was no evidence for ongoing N1575Y selection in either species (□^2^ =0.026; *p*=0.873 and □^2^ =0.62; *p*=0.433 for *An. coluzzii* and *An. gambiae* s.s., respectively). For *An. coluzzii*, there was no significant association between N1575Y frequency and ability of mosquitoes to survive pyrethroid exposure, in LLIN or resistance bioassay (□^2^ =0.0001; *p*=0.993 and □^2^ =0.3244; *p*=0.569, respectively).

G119S *Ace-1* screening revealed 55.1% (27/49) of *An. gambiae* s.l. mosquitoes harboured the mutation; all of these were heterozygotes. G119S *Ace-1* prevalence was 64.9% (24/37) and 27.3% (3/11) for PCR-confirmed *An. coluzzii* and *An. gambiae* s.s., respectively; one remaining *An. gambiae*-*An. coluzzii* hybrid was wild type. For *An. coluzzii*, population-level G119S *Ace-1* allele frequency was 0.32, with evidence for significant deviations from Hardy-Weinberg equilibrium (□^2^ =8.525; *p*=0.00350). For *An. gambiae* s.s., population-level G119S *Ace-1* allele frequency was 0.14, with no significant deviations from Hardy-Weinberg equilibrium (□^2^ =0.274; *p*=0.6005). For *An. coluzzii*, there was a significant association between presence of the G119S *Ace-1* mutation and surviving bendiocarb exposure (Fisher’s exact test = 0.005).

### Metabolic resistance mechanisms

Comparison of metabolic gene expression levels in field populations of *An. coluzzii* and *An. gambiae* s.s. demonstrated significant upregulation of *CYP6P4* (FC=5.88, 95% CI: 5.19-44.06; and 6.08, 95% CI: 5.43-50.64), *CPY6Z1* (FC=4.04, 95% CI: 3.69-41.54; and 3.56, 95% CI: 3.24-36.25) and *CYP6P3* (FC=12.56, 95% CI: 11.40-123.83; and 13.85, 95% CI: 12.53-132.03), relative to a susceptible laboratory colony, respectively (Figure 4). More modest overexpression of *CYP6P1* and *GSTE2* was observed (FC=1.18, 95% CI: 1.08-12.31; and 1.28, 95% CI: 1.17-14.40; FC=0.56, 95% CI: 0.48-3.32; and 0.67, 95% CI: 0.58-4.29; for *An. coluzzii* and *An. gambiae* s.s., respectively) (Figure 4). Levels of FC did not differ significantly between the two species for any gene nor by malaria infection status in wild *An. coluzzii*.

**Figure 4.**
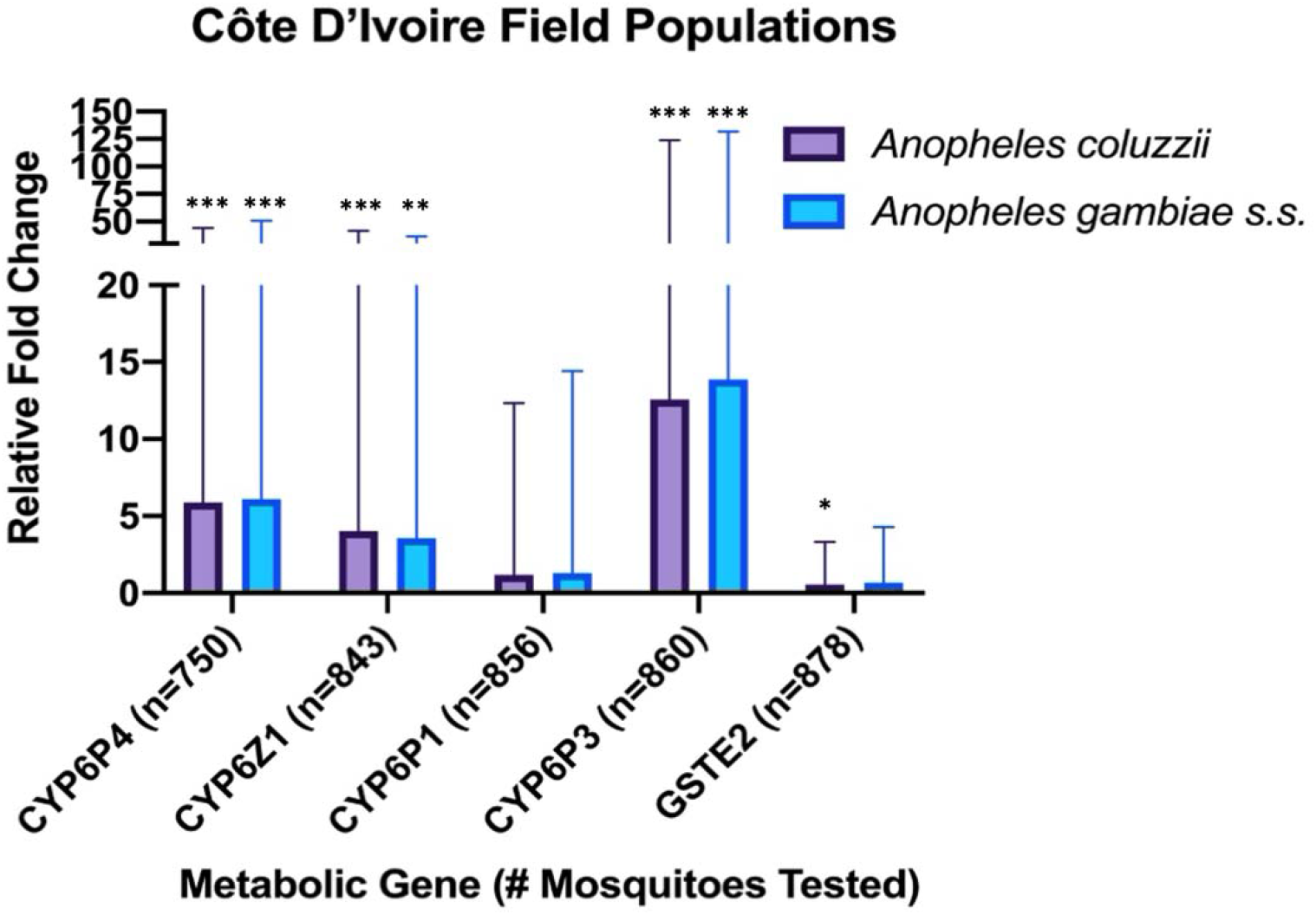
Metabolic gene expression in field *An. coluzzii* and *An. gambiae* s.s. populations relative to a susceptible colony population. Error bars represent 95% CI; statistically significant differences in expression levels relative to the susceptible colony are indicated as \**P*<0.05, \*\**P*<0.01, \*\*\**P*≤0.001.

Comparison of metabolic gene expression in phenotyped field populations of *An. coluzzii* revealed lower FCs overall, but notably, increased overexpression of *CYP6P3* in survivors of bendiocarb, deltamethrin, PBO + deltamethrin and permethrin (FC = 3.91, 95% CI: 3.33-22.16; 2.21, 95% CI: 1.88-12.53; 2.64, 95% CI: 2.21-13.69; and 2.21, 95% CI: 1.99-20.03, respectively) (Figure 5).

**Figure 5.**
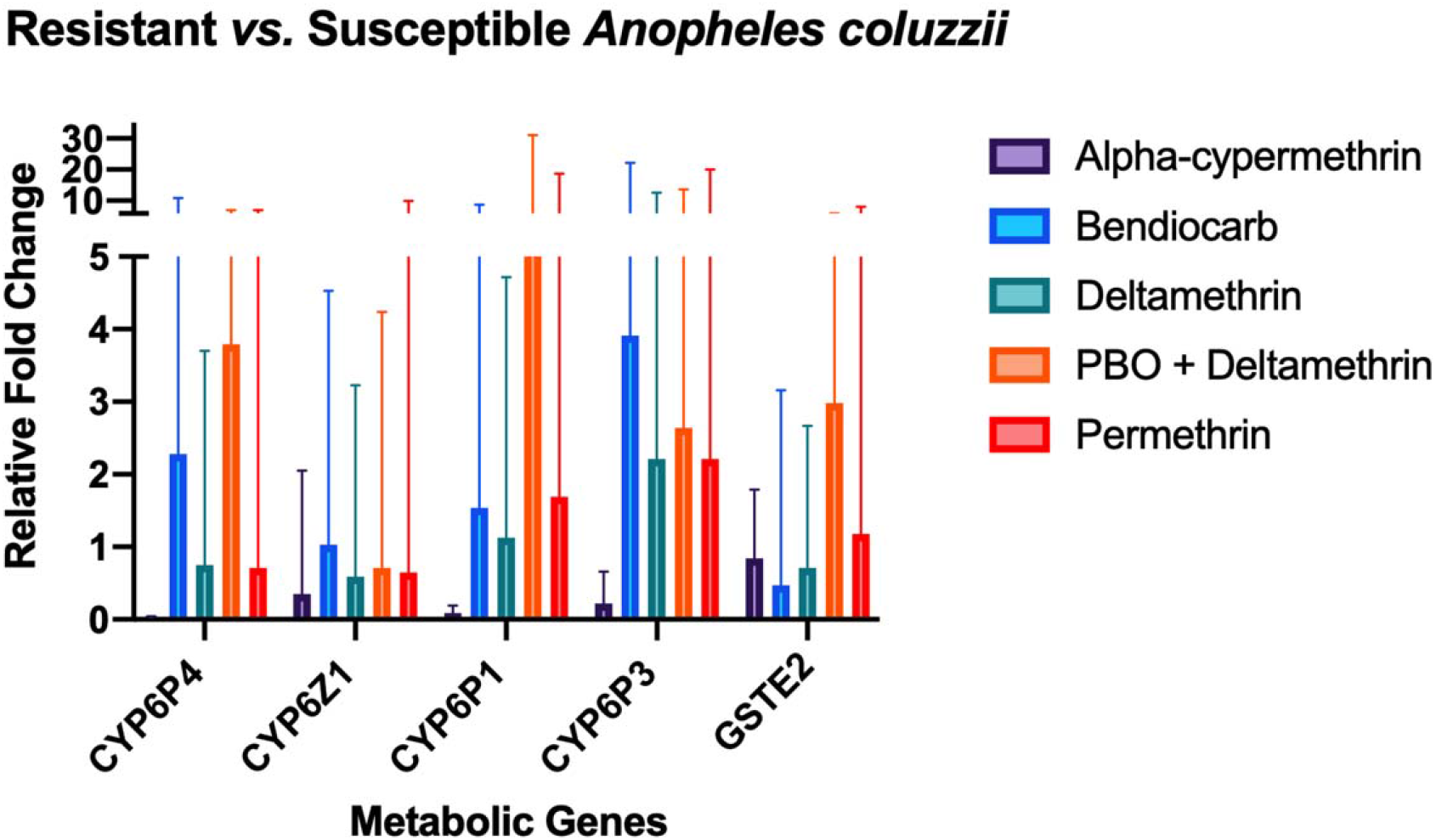
Metabolic gene expression in resistant *versus* susceptible field *An. coluzzii*, which either died or survived following insecticidal exposure. Error bars represent 95% CI.

## Discussion

Côte d’Ivoire is a hot spot of some of the highest levels of resistance of *Anopheles* mosquitoes to public health insecticides worldwide, with potentially severe implications for sustaining gains in malaria control [31]. To safeguard future malaria vector control efforts and inform the design of effective resistance management strategies, involving tactical deployment of differing IRS and LLIN modalities, there needs to be a clear understanding of contemporary phenotypic and genotypic insecticide resistance.

Our study detected intense pyrethroid resistance in south-east, Côte d’Ivoire, as evidenced by high proportions of survivors, following exposure to ten times the diagnostic doses of pyrethroids, as well as very low levels of knock-down and 24-hour mortality to deltamethrin-only LLINs, equivalent to an untreated net. These findings are largely in agreement with historical resistance profiles from this region [7,10,11] and indicate that conventional LLINs may no longer be operationally viable in areas of high pyrethroid resistance intensity. Previous Phase II studies of pyrethroid-only LLINs in the central region of Côte d’Ivoire have demonstrated similarly poor efficacy with highly resistant *An. gambiae* s.l. populations but argued for the retention of some degree of personal protection [15-17]. Other observational cohorts have reported higher incidences of malaria among non-net users compared to users in areas of moderate to high pyrethroid resistance [17]. The extent of protective efficacy afforded by pyrethroid LLINs will likely reflect the strength of local vector resistance and levels of both net physical integrity and individual compliance [32,33]. However, in Côte d’Ivoire, reported LLIN usage has been low, requiring additional behavioural interventions [2,34]. Our findings of high mosquito mortality following exposure to clothianidin and chlorfenapyr and improved vector susceptibility with PBO treatment (on both LLINs and in resistance bioassays), are consistent with data from other sentinel sites across Côte d’Ivoire [16,35,36], and strongly support the deployment of vector control interventions incorporating these new active ingredients.

Study results indicate that *An. coluzzii* was the predominant local vector species during the rainy season, as observed previously [7], circulating sympatrically with smaller proportions of *An. gambiae* s.s.. These two vector species commonly co-habit but can be genetically distinct in terms of resistance mechanisms [37,38] and can also differ in larval ecology, behaviour, migration and aestivation [39-41]. In general, resistance mechanisms in *An. coluzzii* are less well-characterized, compared to *An. gambiae* s.s., in part because these vectors are morphologically indistinguishable and few studies present data disaggregated by PCR-confirmed species. We observed several distinct features in our study, principally, evidence for ongoing selection of L1014F *kdr* and G119S *Ace-1* in *An. coluzzii*, which was absent in *An. gambiae* s.s. and higher proportions of N1575Y in *An. gambiae* s.s.; expression levels of metabolic genes were comparable between species. The lack of association between L1014F *kdr* genotype and mosquito phenotype, coupled with the identification of three CYP450 enzymes (*CYP6P4, CYP6P3* and *CYP6Z1*) that were significantly over-expressed in field populations, (some of which are known to metabolise pyrethroids and next generation LLIN insecticides [42,43]), indicate a key role for metabolic resistance in this *An. coluzzii* population. One notable difference in our dataset, compared to previous work in Agboville [7], was the finding of bendiocarb susceptibility. This may be attributable to small-scale spatial and longitudinal heterogeneity in resistance, which can be highly dynamic [37,44], and/or phenotypic differences between vector species.

With the exception of chlorfenapyr, which is known to be a slow-acting insecticide, we did not detect any delayed mortality effects for 72 hours following insecticidal exposure; the format and dose used for clothianidin testing (another slow-acting insecticide [45]) was instead intended to measure acute toxicity within a 60 minute exposure period. Previous mathematical models using resistant mosquito colonies have suggested that sub-lethal insecticide treatment may still reduce vector lifespan and inhibit blood-feeding and host-seeking behaviours, thereby interrupting malaria transmission [46,47]. Our observations are more compatible with reports from Burkina Faso where different exposure regimens of wild, resistant *An. gambiae* s.l. populations to deltamethrin LLINs did not induce any delayed mortality [47]. Further assessment of sublethal effects are warranted across additional field populations with differing resistance mechanisms to better understand the impact of insecticidal exposure on vectorial capacity of resistant mosquitoes.

To date there is a paucity of data regarding the interactions between insecticide resistance and *Plasmodium* development [48]. In this study, *An. coluzzii* which died following pyrethroid exposure were significantly more likely to be infected with malaria. This might be explained by elevated metabolic enzymes and/or prior pyrethroid exposure detrimentally affecting parasite development [49]; although it is important to note that we did not detect any significant differences between gene overexpression in malaria infected *vs*. non-infected *An. coluzzii*. Alternatively, our sampled population may have been physiologically older, as phenotypic resistance is known to decline with age [50]. It is impossible to distinguish between these hypotheses using field-collected vector populations; the experimental design used in this study had other biological and technical limitations, which have been described in detail previously [23,37].

## Conclusions

As new combination and bi-treated vector control interventions become available for deployment, contemporary resistance information is crucial for the rationale design of management strategies and to mitigate future selection for particular resistance mechanisms. The results from this study contribute to growing insecticide resistance data for Côte d’Ivoire, demonstrating a loss of bioefficacy of conventional pyrethroid LLINs and supporting the use of new active ingredients (clothianidin, chlorfenapyr and PBO). Study findings also highlight the need for expanded insecticide resistance surveillance, including monitoring of metabolic resistance mechanisms, in conjunction with studies to better characterize the impact of sublethal insecticide exposure on vectorial capacity and the interaction between insecticide resistance on *Plasmodium* parasite development.

## Acknowledgements

The authors express their sincere thanks to the M. Didier Dobri, CSRS lab technician, and Fidele Assamoa for their support in mosquito collection and rearing, the chief and population of the village of Aboudé (Agboville) and the entomology fieldworkers of CSRS. Study funding was provided by a Sir Halley Stewart Trust (awarded to LAM) and a Wellcome Trust/Royal Society grant (awarded to TW; 101285/Z/13/Z) http://www.wellcome.ac.uk; https://royalsociety.org. SI is supported by the President’s Malaria Initiative (PMI)/CDC. The findings and conclusions in this report are those of the author(s) and do not necessarily represent the official position of the Centers for Disease Control and Prevention.

## Author contributions

AM, EM, MK, TW and LAM designed the study. AM, EM, CE and BP led the entomology field activities and participated in data collection. AM, EM, CLJ, TW and LAM performed the molecular assays. AM, EM, MK, CE, CLJ, BP, SI, TW and LAM were responsible for data analysis and interpretation. LAM drafted the manuscript, which was revised by all co-authors. All authors read and approved the final manuscript.

## Conflict of interest

The authors declare no conflict of interest.

## Notes

### Competing Interest Statement

The authors have declared no competing interest.

## References

1. Institut National de la Statistique (INS), Programme National de Lutte contre le Paludisme (PNLP) et ICF. 2016. Enquête de prévalence parasitaire du paludisme et de l’anémie en Côte d’Ivoire 2016. Rockville, Maryland, USA : INS, PNLP et ICF.

2. USAID. President’s Malaria Initiative: Côte d’Ivoire: Malaria Operational Plan FY 2018 and FY 2019, USAID; 2019. Available from: https://www.pmi.gov/docs/default-source/default-document-library/malaria-operational-plans/fy19/fy-2019-cote-d'ivoire-malaria-operational-plan.pdf?sfvrsn=5.

3. Killeen GF, Ranson H. Insecticide-resistant malaria vectors must be tackled. Lancet 2018; 391(10130):1551–1552.

4. Elissa N, Mouchet J, Riviere F, Meunier JY, Yao K. Resistance of *Anopheles gambiae* s.s. To pyrethroids in Côte D’Ivoire. Ann Soc Belge Med Trop 1993; 73:291–294.

5. Elissa N, Mouchet J, Riviere F, Meunier JY, Yao K. Sensibilite d’*Anopheles gambiae* aux insecticides en Côte D’Ivoire. Cahiers Sante 1994; 4:95–99.

6. N’Guessan R, Darriet F, Guillet P, et al. Resistance to carbosulfan in *Anopheles gambiae* from Ivory Coast, based on reduced sensitivity of acetylcholinesterase. Med Vet Entomol 2003; 17:19–25.

7. Edi CAV, Koudou BG, Bellai L, et al. Long-term trends in *Anopheles gambiae* insecticide resistance in Côte D’Ivoire. Parasit Vectors 2014; 7:500.

8. Chandre F, Darriet F, Manga L, Akogbeto M, Faye O, Mouchet J, Guillet P. Status of pyrethroid resistance in *Anopheles gambiae* sensu lato. Bull World Health Organ 1999; 77:230–234.

9. Koffi AA, Alou LP, Adja MA, Kone M, Chandre F, N’Guessan R. Update on resistance status of *Anopheles gambiae* s.s. To conventional insecticides at a previous WHOPES field site, “Yaokoffikro”, 6 years after the political crisis in Côte D’Ivoire. Parasit Vectors 2012; 5:68.

10. Edi CVA, Koudou BG, Jones CM, Weetman D, Ranson H. Multiple-insecticide resistance in *Anopheles gambiae* mosquitoes, Southern Côte D’Ivoire. Emerg Infect Dis 2012; 18:1508–1511.

11. Camara S, Koffi AA, Alou LPA, et al. Mapping insecticide resistance in *Anopheles gambiae* (s.l.) from Côte D’Ivoire. Parasit Vectors 2018; 11:19.

12. Mouhamadou CS, de Souza SS, Fodjo BK, Zoh MG, Bli NK, Koudou BG. Evidence of insecticide resistance selection in wild *Anopheles coluzzii* mosquitoes due to agricultural pesticide use. Infect Dis Poverty 2019; 8:64.

13. Fodjo BK, Koudou BG, Tia E, et al. Insecticides Resistance Status of *An. gambiae* in Areas of Varying Agrochemical Use in Côte D’Ivoire. Biomed Res Int 2018; 2018:2874160.

14. Chouaibou MS, Fodjo BK, Fokou G, et al. Influence of the agrochemicals used for rice and vegetable cultivation on insecticide resistance in malaria vectors in southern Côte d’Ivoire. Malar J 2016; 15:426.

15. Oumbouke WA, Koffi AA, Alou LPA, Rowland M, N’Guessan R. Evaluation of standard pyrethroid based LNs (MiraNet and MagNet) in experimental huts against pyrethroid resistance *Anopheles gambiae* s.l. M’be, Côte D’Ivoire: potential for impact on vectorial capacity. PLoS One 2019; 14(4):e0215074.

16. Oumbouke WA, Rowland M, Koffi AA, Alou LPA, Camara S, N’Guessan R. Evaluation of an alpha-cypermethrin + PBO mixture long-lasting insecticidal net VEERALIN LN against pyrethroid resistant *Anopheles gambiae* s.s.: an experimental hut trial in M’be, central Côte D’Ivoire. Parasit Vectors 2019; 12:544.

17. Kleinschmidt I, Bradley J, Knox TB, et al. Implications of insecticide resistance for malaria vector control with long-lasting insecticidal nets: a WHO-coordinated, prospective, international, observational cohort study. Lancet Infect Dis 2018; 18:640–649.

18. Edi AVC, N’Dri BP, Chouaibou M, et al. First detection of N1575Y mutation in pyrethroid resistant *Anopheles gambiae* in Southern Côte D’Ivoire. Wellcome Open Res 2017; 2:71.

19. Edi CV, Djogbenou L, Jenkins AM, et al. CYP6 P450 enzymes and *ACE-1* duplication produce extreme and multiple insecticide resistance in the malaria mosquito *Anopheles gambiae*. PLoS Genet 2014; 10(3):e1004236.

20. Gillies, M. T. & Coetzee, M. A supplement of the Anophelinae of Africa south of the Sahara (Afrotropical Region). Publ. South African Inst. Med. Res. No. 55 (1987).

21. World Health Organisation (WHO). Guidelines for Laboratory and Field-Testing of Long-lasting Insecticidal Nets. 2013.

22. Centers for Disease Control and Prevention. Guideline for Evaluating Insecticide Resistance in Vectors Using the CDC Bottle Bioassay. CDC Methods 1–28 (2012).

23. Stica C, Jeffries CL, Irish SR, et al. Characterizing the molecular and metabolic mechanisms of insecticide resistance in Anopheles gambiae in Faranah, Guinea. Malar J 2019; 18(1):244.

24. World Health Organization. Test procedures for insecticide resistance monitoring in malaria vector mosquitoes. Second edition. https://apps.who.int/iris/bitstream/handle/10665/250677/9789241511575-eng.pdf?sequence=1 (2016.

25. Santolamazza F, Mancini E, Simard F, Qi Y, Tu Z, della Torre A. Insertion polymorphisms of SINE200 retrotransposons within speciation islands of *Anopheles gambiae* molecular forms. Malar J 2008; 7:163.

26. Boissiere A, Gimonneau G, Tchioffo MT et al. Application of a qPCR assay in the investigation of susceptibility to malaria infection of the M and S molecular forms of *An. gambiae s.s.* in Cameroon. PLoS One 2013; 8:e54820.

27. MR4. Methods in Anopheles research (2nd ed.). 2016. Retrieved from https://www.beiresources.org/Portals/2/VectorResources/2016%20Methods%20in%20Anopheles%20Research%20full%20manual.pdf

28. Jones CM, Liyanapathirana M, Agossa FR, Weetman D, Ranson H, Donnelly MJ, Wilding CS. Footprints of positive selection associated with a mutation (N1575Y) in the voltage-gated sodium channel of *Anopheles gambiae*. Proc Natl Acad Sci USA 2012; 109:6614–6619.

29. Weill M, Malcolm C, Chandre F, Mogensen K, Berthomieu A, Marquine M, Raymond M. The unique mutation in ace-1 giving high insecticide resistance is easily detectable in mosquito vectors. Insect Mol Biol 2004; 13:1–7.

30. Mavridis K, Wipf N, Medves S, Erquiaga I, Muller P, Vontas J. Rapid multiplex gene expression assays for monitoring metabolic resistance in the major malaria vector *Anopheles gambiae*. Parasit Vectors 2019; 12:9.

31. Glunt KD, Coetzee M, Huijben S, et al. Empirical and theoretical investigation into the potential impacts of insecticide resistance on the effectiveness of insecticide-treated bed nets. Evol Appl. 2018; 11:431–441.

32. Shah MP, Steinhardt LC, Mwandama D, et al. The effectiveness of older insecticide-treated bed nets (ITNs) to prevent malaria infection in an area of moderate pyrethroid resistance: results from a cohort study in Malawi. Malar J 2020; 19:24.

33. Toe KH, Jones CM, N’Fale S, Ismail HM, Dabire RK, Ranson H. Increased pyrethroid resistance in malaria vectors and decreased bed net effectiveness, Burkina Faso. Emerg Infect Dis 2014; 20(10):1691–1696.

34. Ouattara AF, Raso G, Edi CVA, Utzinger J, Tanner M, Dagnogo M, Koudou BG. Malaria knowledge and long-lasting insecticidal net use in rural communities of central Côte d’Ivoire. Malar J 2011; 10:288.

35. Kouassi BL, Edi C, Tia E, et al. Susceptibility of An. gambiae s.l. from Côte d’Ivoire to insecticides used on insecticide-treated nets: evaluating the additional entomological impact of piperonyl butoxide and chlorfenapyr. PrePrint, accessed from https://www.researchsquare.com/article/rs-21138/v1.

36. Camara S, Ahoua Alou LP, Koffi AA, et al. Efficacy of Interceptor®G2, a new long-lasting insecticidal net against wild pyrethroid-resistant *Anopheles gambiae* s.s. From Côte d’Ivoire: a semi-field trial. Parasite 2018; 25:42.

37. Collins E, Vaselli NM, Sylla M, et al. The relationship between insecticide resistance, mosquito age and malaria prevalence in *Anopheles gambiae* s.l. from Guinea. Sci Rep 2019; 9:8846.

38. The Anopheles gambiae 1000 Genome Consortium. Genetic diversity of the African malaria vector *Anopheles gambiae*. Nature 2017; 552(7683):96–100.

39. Fossog BT, Ayala D, Acevedo P, et al. Habitat segregation and ecological character displacement in cryptic African malaria mosquitoes. Evol Appl 2015; 8(4):326–345.

40. Gimonneau G, Bouyer J, Morand S, Besansky NJ, Diabate A, Simard F. A behavioral mechanism underlying ecological divergence in the malaria mosquito *Anopheles gambiae*. Behav Ecol 2010; 21(5):1087–1092.

41. Dao A, Yaro AS, Diallo M, et al. Signatures of aestivation and migration in Sahelian malaria mosquito populations. Nature 2014; 516:387–390.

42. Muller P, Wart E, Stevenson BJ, et al. Field-caught permethrin-resistant *Anopheles gambiae* overexpress CYP6P3, a P450 that metabolises pyrethroids. PLoS Genet 2008; 4(11):e1000286.

43. Yunta C, Grisales N, Nasz S, et al. Pyriproxyfen is metabolised by P450s associated with pyrethroid resistance in *An. gambiae*. Insect Biochem Mol Biol 2016; 78:50–57.

44. Implications of Insecticide Resistance Consortium. Implications of insecticide resistance for malaria vector control with long-lasting insecticidal nets: trends in pyrethroid resistance during a WHO-coordinated multi-country prospective study. Parasit Vectors 2018; 11(1):550.

45. Oxborough RM, Seyoum A, Yihdego Y, et al. Susceptibility testing of *Anopheles* malaria vectors with the neonicotinoid insecticide clothianidin; results from 16 African countries, in preparation for indoor residual spraying with new insecticide formulations. Malar J 2019; 18(1):264.

46. Viana M, Hughes A, Matthiopoulos J, Ranson H, Ferguson HM. Delayed mortality effects cut the malaria transmission potential of insecticide-resistant mosquitoes. Proc Natl Acad Sci USA 2016; 113(32):8975–8980.

47. Hughes A, Lissenden N, Viana M, Toe KH, Ranson H. *Anopheles gambiae* populations from Burkina Faso show minimal delayed mortality after exposure to insecticide-treated nets. Parasit Vectors 2020; 13:17.

48. Minetti C, Ingham VA, Ranson H. Effects of insecticide resistance and exposure on Plasmodium development in *Anopheles* mosquitoes. Curr Opin Insect Sci 2020; 39:42–49.

49. Kristan M, Lines J, Nuwa A, Ntege C, Meek SR, Abeku TA. Exposure to deltamethrin affects development of *Plasmodium falciparum* inside wild pyrethroid resistant *Anopheles gambiae* s.s. mosquitoes in Uganda. Parasit Vectors 2016; 24(9):100.

50. Jones CM, Sanou A, Guelbeogo WM, Sagnon NF, Johnson PCD, Ranson H. Aging partially restores the efficacy of malaria vector control in insecticide-resistant populations of *Anopheles gambiae* s.l. from Burkina Faso. Malar J 2012; 11:24.

